# Rapid Isothermal Amplification and Portable Detection System for SARS-CoV-2

**DOI:** 10.1101/2020.05.21.108381

**Authors:** A. Ganguli, A. Mostafa, J. Berger, M. Aydin, F. Sun, E. Valera, B. T. Cunningham, W. P. King, R. Bashir

## Abstract

The COVID-19 pandemic provides an urgent example where a gap exists between availability of state-of-the-art diagnostics and current needs. As assay details and primer sequences become widely known, many laboratories could perform diagnostic tests using methods such as RT-PCR or isothermal RT-LAMP amplification. A key advantage of RT-LAMP based approaches compared to RT-PCR is that RT-LAMP is known to be robust in detecting targets from unprocessed samples. In addition, RT-LAMP assays are performed at a constant temperature enabling speed, simplicity, and point-of-use testing. Here, we provide the details of an RT-LAMP isothermal assay for the detection of SARS-CoV-2 virus with performance comparable to currently approved tests using RT-PCR. We characterize the assay by introducing swabs in virus spiked synthetic nasal fluids, moving the swab to viral transport medium (VTM), and using a volume of that VTM for performing the amplification without an RNA extraction kit. The assay has a Limit-of-Detection (LOD) of 50 RNA copies/μL in the VTM solution within 20 minutes, and LOD of 5000 RNA copies/μL in the nasal solution. Additionally, we show the utility of this assay for real-time point-of-use testing by demonstrating detection of SARS-CoV-2 virus in less than 40 minutes using an additively manufactured cartridge and a smartphone-based reader. Finally, we explore the speed and cost advantages by comparing the required resources and workflows with RT-PCR. This work could accelerate the development and availability of SARS-CoV-2 diagnostics by proving alternatives to conventional laboratory benchtop tests.

**Significance Statement:** An important limitation of the current assays for the detection of SARS-CoV-2 stem from their reliance on time- and labor-intensive and laboratory-based protocols for viral isolation, lysis, and removal of inhibiting materials. While RT-PCR remains the gold standard for performing clinical diagnostics to amplify the RNA sequences, there is an urgent need for alternative portable platforms that can provide rapid and accurate diagnosis, potentially at the point-of-use. Here, we present the details of an isothermal amplification-based detection of SARS-CoV-2, including the demonstration of a smartphone-based point-of-care device that can be used at the point of sample collection.

## Introduction

Since the coronavirus 2 (SARS-CoV-2) jumped from an animal reservoir to humans in December, 2019, the acute respiratory disease (COVID-19) has rapidly spread across the world, bringing death, illness, disruption to daily life, and economic losses to businesses and individuals^1–3^. The rapid development of the COVID-19 pandemic highlights shortcomings in the existing laboratory-based testing paradigm for viral diagnostics^4^. The fundamental limitations of current diagnostics assays for viral pathogens stem from their reliance upon polymerase chain reaction (PCR) analysis, which requires labor-intensive, laboratory-based protocols for viral isolation, lysis, and removal of inhibiting materials. While PCR remains the proven gold standard for clinical diagnostics, there is an urgent need for other approaches that are low-cost, rapid and provide diagnosis at the point of use.

In addition to the CDC RT-PCR SARS-CoV-2 test^5^, other diagnostic tests have become available including the Cepheid Xpert® Xpress SARS-CoV-2 test^6^, Abbott ID NOW™ COVID-19 test^7^, and others^8–18^. The Cepheid SARS-CoV-2 test can provide results for the detection of SARS-CoV-2 in approximately 45 minutes^19^ with low false-negative rate at 1.8% demonstrated in an independent study^20^. However, this test requires the GeneXpert system, of which there are only 5,000 systems available in the US^21^. The test also requires RNA extraction as a separate step from amplification and detection, which is a key constraint on scalability and could become important as demand increases for critical supplies^22^. The Abbott ID NOW isothermal amplification technology claims the delivery of positive results in less than 15 minutes while offering a device with portable size and weight. However, this test also requires a specialized instrument and availability issues^23–25^ as well as accuracy issues have been recently reported for this test^20^. The root causes of accuracy problems are unknown and this issue is further being evaluated. At the time of submission this paper, the events and information are rapidly changing around this topic.

While the current laboratory-based paradigm for SARS-CoV-2 should be scaled as quickly as possible, there is an urgent need for alternatives to current approaches to further expand the options for testing. Laboratory tests require expensive capital equipment, laboratory infrastructure, and human resources with specialized expertise. While these resources are normally available in densely populated and wealthy regions of the world, much of the world lacks one or more of these elements. Commercially available COVID-19 diagnostic tests in the U.S, Europe or Asia are generally benchtop type systems and are not tailored for portability and point-of-use applications^26^. To broaden access to testing, there is a need for technologies that are fast, low cost, can be performed away from the laboratory without specialized capital equipment, and that require minimal training and expertise.

In recent years, LAMP amplification has emerged as a compelling alternative to PCR, because LAMP can be performed without the need for commercial thermocyclers^27^. LAMP-based diagnostics can be faster than PCR because LAMP does not require time for thermal cycling, compared to PCR, which requires time for the thermal cycler to progress through different temperatures. The simplicity of isothermal amplification also allows for translation to simple point-of-use devices that use disposable cartridges^28^. RT-LAMP has advantages over RT-PCR for targeting sequences due to its robustness against inhibitors^29,30^ as well as its high specificity using 4-6 primers that identify 6-8 regions on the template for amplification^27^.

This paper presents detailed characterization of a SARS-CoV-2 diagnostic test based on an RT-LAMP assay using viral transport media and synthetic spiked nasal solutions. We also demonstrate the potential of point-of-use detection using an additively manufactured three-dimensional cartridge and a smartphone-based optical measurement. We present an analysis of the resources required to scale up the assay. The proposed approach avoids the necessity of RNA extraction and could impact the diagnosis of the COVID-19 outbreak specially in settings where diagnosis is required at the point of collection or regions that lack laboratory-grade infrastructure and resources.

## Results

### Primer design and assay characterization with SARS-CoV-2 genomic RNA

To develop a sensitive and specific RT-LAMP assay for the detection of SARS-CoV-2 virus, we designed sequence-specific primers for four genes from the SARS-CoV-2 viral genome. Using BLASTn analysis, we identified genes Orf 1a, S, Orf 8, and N for primer design, which code for the orf1ab polyprotein, surface glycoprotein, Orf 8 protein, and nucleocapsid phosphoprotein, respectively (**Figure 1a**). Target regions Orf 1ab, S, and Orf 8 were selected because they showed the least similarity with other Coronavirus sequences such as SARS-CoV-1 and MERS-CoV^31^. Gene N target region was selected due to its overlap with the region used for primer design in currently CDC and FDA approved assays for COVID19^32^. Three primer sets for each of the four selected genes were generated using Primerexplorerv4 (https://primerexplorer.jp/e/), and RT-LAMP experiments using SARS-CoV-2 genomic RNA were performed in a standard thermocycler. **Figure 1b-e** show the amplification curves for the detection of 500 copies/uL of SARS-CoV-2 genomic RNA using primer sets for Gene Orf 1a, Gene S, Gene Orf 8, and Gene N, respectively. **Figure 1f** shows the threshold times for detection for all the above primer sets. The best primer set for each gene (primer set 3 for Gene Orf 1a, primer set 2 for Gene S, primer set 2 for Gene Orf 8, and primer set 1 for Gene N) was selected based on lowest threshold time and used for limit of detection analysis. All the primer sequences tested are shown in **Table S1** (Supplementary information).

**Fig. 1.**
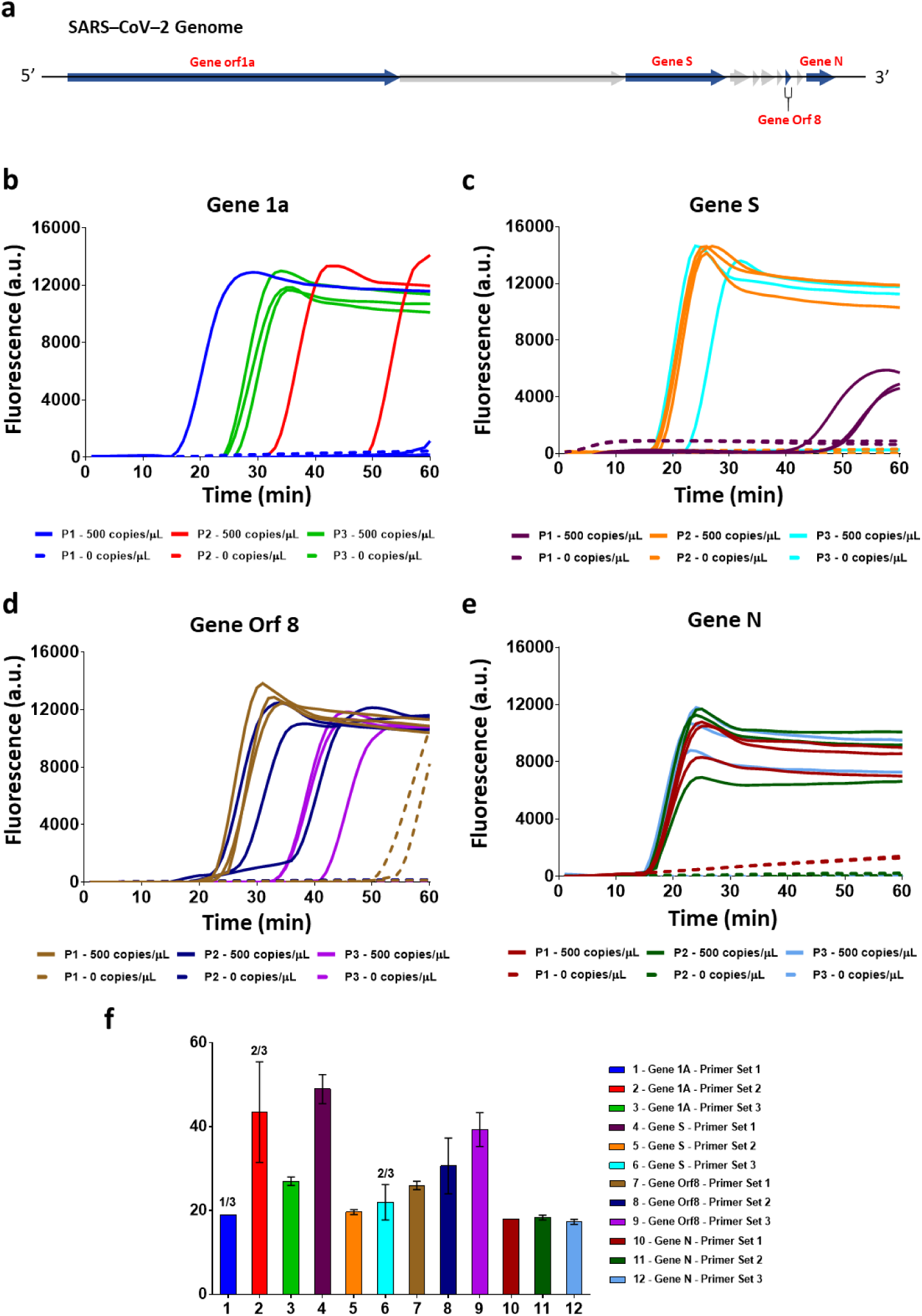
Validation of 3 LAMP primer sets for 4 different SARS-CoV-2 Gene Targets. **(*a*)** SARS-CoV-2 genome outline and 4 genes targets for primer design. **(*b-e*)** Raw fluorescence data for detecting Gene orf 1a **(*b*)**, Gene S **(*c*)**, Gene orf 8 **(*d*)**, Gene N **(*e*)** of SARS-COV-2 using three different primer sets for each gene. **(*f*)** Comparison of positive amplification threshold time for four genes from 3 replicates of amplification data seen in Fig. 1b-e. The bar graphs show mean and standard deviation.

Next, we compared the limit of detection of the four selected primers sets by amplifying serial dilutions of SARS-CoV-2 RNA (**Figure 2**). The limit of detection for RNA using the Gene Orf 1a primer set was 500 copies/μL with only 2/3 replicates giving amplification for 50 copies/μL of sample (**Figure 2a-b**). **Figure 2c-f** demonstrates a detection limit of 5000 copies/μL of RNA using the Gene S and Gene Orf 8 primers, respectively, as not all replicates amplified for 500 copies/μL of sample. The reactions performed with Gene N primers demonstrated the lowest limit of detection and fastest amplification times with 50 copies/μL amplifying within 25 min. of the reaction start (**Figure 2g-h**). Hence, we chose primer set 1 targeting Gene N as the final primer set for our RT-LAMP assay for SARS-CoV-2 detection.

**Fig. 2.**
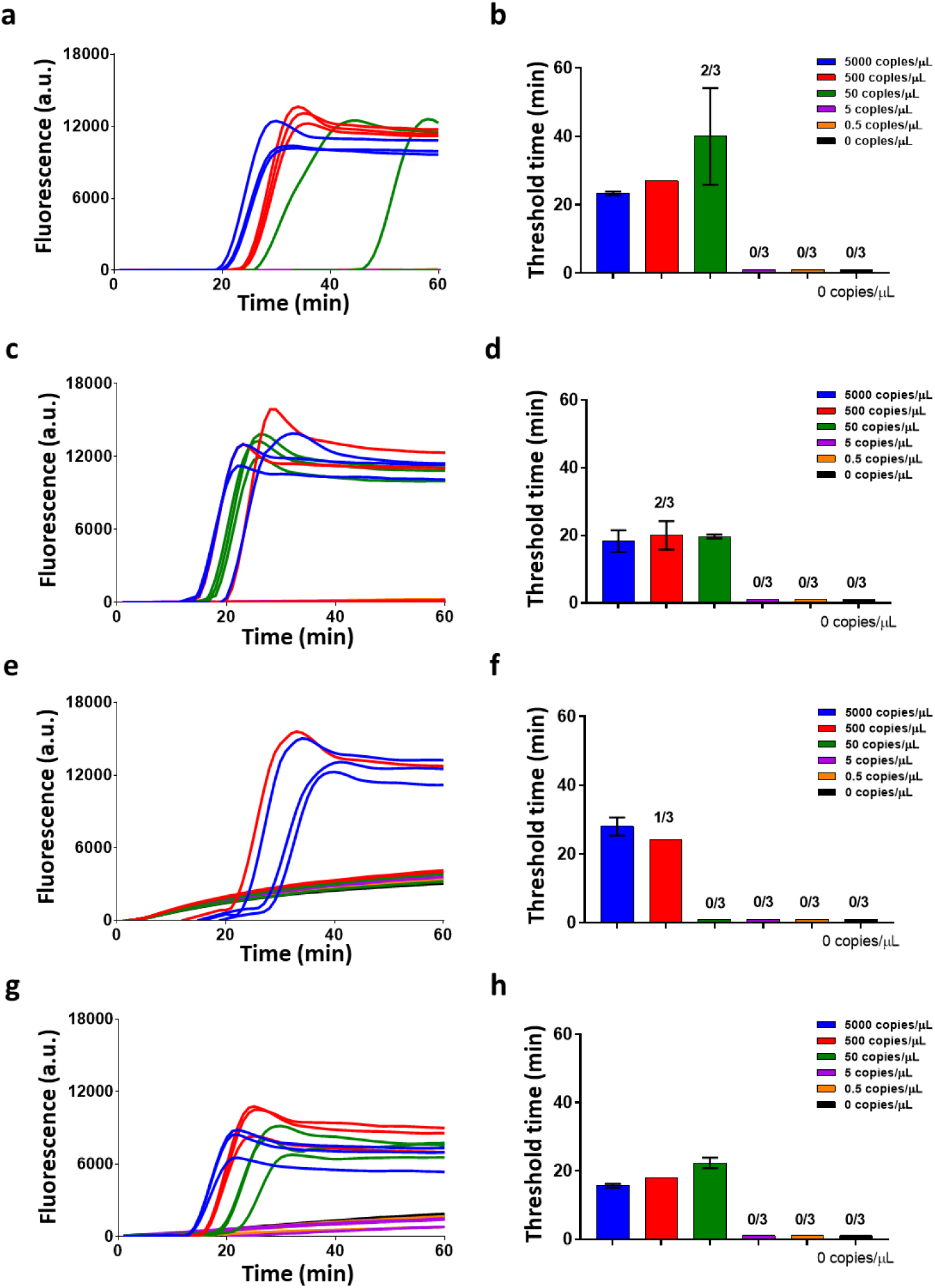
Characterization of SARS-CoV-2 genomic RNA in buffer. **(*a-h*)** Raw fluorescence data and amplification threshold times (for 3 replicates of data) for detection of genomic RNA using primer set 3 for Gene orf 1a **(*a-b*)**, primer set 2 for Gene S **(*c-d*)**, primer set 2 for Gene orf 8 **(*e-f*)**, and primer set 1 for Gene N **(*g-h*)**. The bar graphs show mean and standard deviation. The best detection limit was 50 copies/uL attained using Gene N primer set 1.

### Assay characterization with SARS-CoV-2 inactive viruses

To evaluate the efficacy of our RT-LAMP assay to detect SARS-CoV-2, we next prepared serial dilutions of spiked inactive SARS-CoV-2 viruses in our 16μL RT-LAMP reactions. The amplification curves and threshold times are shown in **Figure 3a-b**. The detection limit was 500 copies/μL (0.005 PFU/μL) of starting virus concentration with only 2/3 replicates amplifying for 50 copies/μL starting concentration. The above reaction did not include any additional viral lysis step, and the reaction was directly performed at 65°C. Next, to evaluate the effect of a short thermal lysis step on the detection limit of our assay, we repeated the above experiment but with an additional step of thermal lysis of the viral sample at 95°C for 1 minute before addition to the RT-LAMP reaction. We see an improvement of 1 order in the LOD of the assay with a detection limit of 50 copies/μL, where all three replicates amplified within 20 minutes of the reaction start **Figure 3c-d**. Thermal lysis not only efficiently disrupts the viral cell membrane to expose the RNA for amplification but has also been shown to inactivate the nucleases that are present in crude and unpurified samples^33^. Henceforth, a 95°C, 1-minute thermal lysis was performed for all viral samples in the following experiments.

**Fig. 3.**
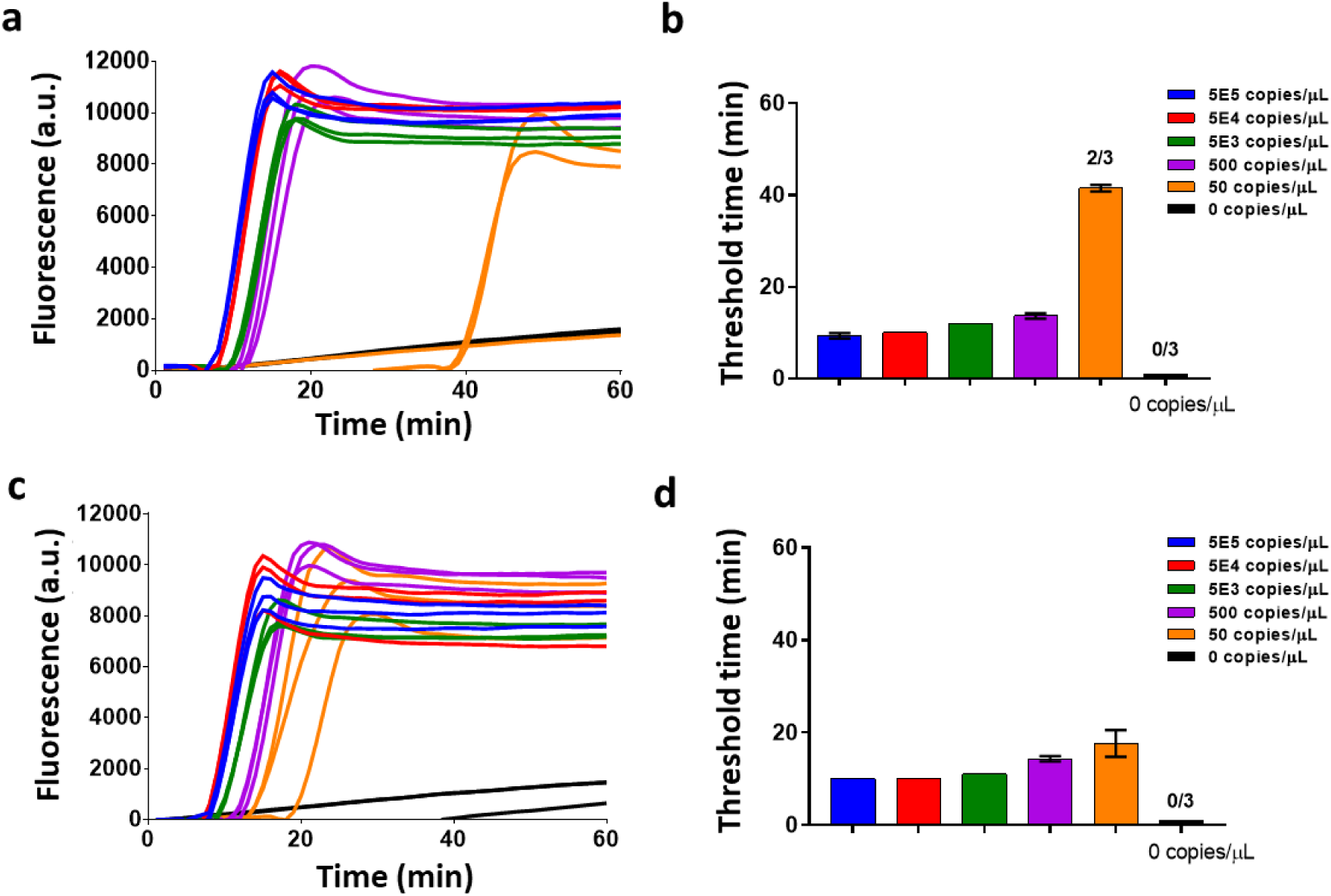
Characterization of thermal lysis of inactive SARS-CoV-2 virus in buffer. **(*a-b*)** Raw fluorescence data and amplification threshold times (for 3 replicates of data) for inactive SARS-CoV-2 virus detection without a thermal lysis step prior to the final reaction. **(*c-d*)** Raw fluorescence data and amplification threshold times (for 3 replicates of data) for inactive SARS-CoV-2 virus detection in a reaction with an additional thermal lysis step conducted at 95°C for 1 min. prior to the addition of RT-LAMP reagents for the final reaction. The bar graphs show mean and standard deviation.

### Assay characterization with SARS-CoV-2 viruses spiked in crude human nasal fluid samples

To characterize the robustness of our RT-LAMP assay, we spiked inactive SARS-CoV-2 viruses directly into purchased healthy human nasal fluid samples. To evaluate the effect of volume percentage of spiked nasal fluid per reaction on the detection limits and amplification times, we varied the spiked nasal sample per reaction from 12.5% to 50% of the total reaction volume. The amplification curves and threshold times of these reactions are shown in **Figures 4a-f.** Surprisingly, we observed that the RT-LAMP reactions showed robust amplifications even for 50% nasal fluid per reaction. Moreover, we observed that higher sampling volumes of virus-spiked nasal fluid improved the detection limit of the assay from 5E5 copies/μL (for 12.5% nasal sample/reaction) to 5E3 copies/μL (for 50% nasal sample/reaction) in the final reaction. This is likely because the very viscous nasal fluid solution prevented effective pipette mixing and caused heterogeneous distribution of the viral target. Thus, higher nasal volumes allowed for sampling more viral particles from the inhomogeneous sample yielding better results. For a 12.5% sample, we sampled only 2 μL of spiked nasal fluid in a 16 μL final reaction, compared to 8 μL nasal fluid added in a 50% sample per reaction. As volume of crude nasal sample increased in the final reaction, there was a delay in amplification times by 3 min. for the 25% sample and by 8 min. for the 50% sample compared to the amplification times of the 12.5% sample for 5.5E5 copies/μL of viral concentration in the nasal sample. As expected, with increasing amounts of crude sample per reaction, the increase in the inhibitory components causes a delay in amplification times. Finally, to evaluate whether sampling more of the spiked nasal fluid in a reaction improves the detection limit, we performed 96 μL reactions, where the spiked nasal fluid sample volume was 48 μL (50% nasal fluid/reaction). **Figure S1** (Supplementary information) shows the amplification curves and threshold times of these reactions, but no improvement in the LOD was seen with higher reaction volumes. Together, these results highlight that our reactions can tolerate up to 50% of crude nasal fluid samples per reaction.

**Fig. 4.**
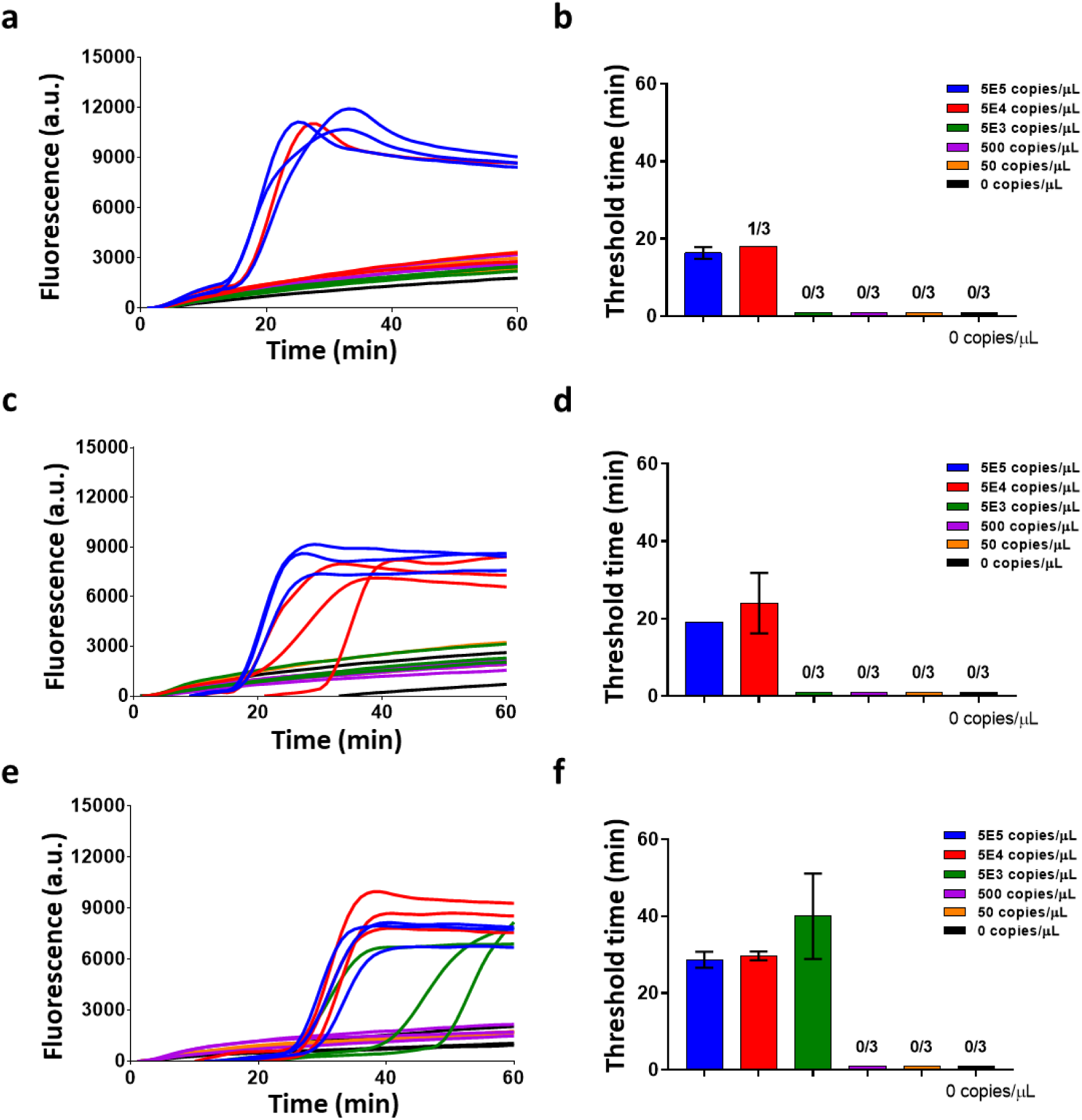
Characterization of SARS-CoV-2 virus in Nasal Fluid. **(*a-b*)** Raw fluorescence data and amplification threshold times (for 3 replicates of data) for viral detection in a 16 μL reaction with 12.5% spiked nasal fluid per reaction. **(*c-d*)** Raw fluorescence data and amplification threshold times (for 3 replicates of data) for viral detection in a 16 μL reaction with 25% spiked nasal fluid per reaction. **(*e-f*)** Raw fluorescence data and amplification threshold times (for 3 replicates of data) for viral detection in a 16 μL reaction with 50% spiked nasal fluid per reaction. The bar graphs show mean and standard deviation.

### Assay characterization with SARS-CoV-2 viruses spiked in Viral Transport Media (VTM)

For diagnostic testing of SARS-CoV-2, the current workflow includes collecting nasopharyngeal/nasal specimens using swabs, which are immediately transferred into sterile transport tube containing 2-3 mL of viral transport media (VTM) until diagnostic assays can be performed^34^. In standard RT-PCR assays, this VTM sample with viruses undergoes an RNA purification step next^32^. In our RT-LAMP assay to evaluate the direct use of VTM samples without any RNA purification, we spiked serial dilutions of inactive SARS-CoV-2 viruses in VTM and performed our RT-LAMP reactions. **Figure 5a-d** show the amplification fluorescence curves and threshold times of reactions where the VTM sample was either 12.5% or 50% of the final reaction volume, and the detection limit for both reactions was 50 copies/μL. These results highlight no loss in the detection limit of our assay in VTM compared to SARS-CoV-2 viruses in buffer. The reactions with 50% VTM showed a delay of ∼20 min. compared to the 12.5%, likely due to the increase in inhibitory components in the VTM.

**Fig. 5.**
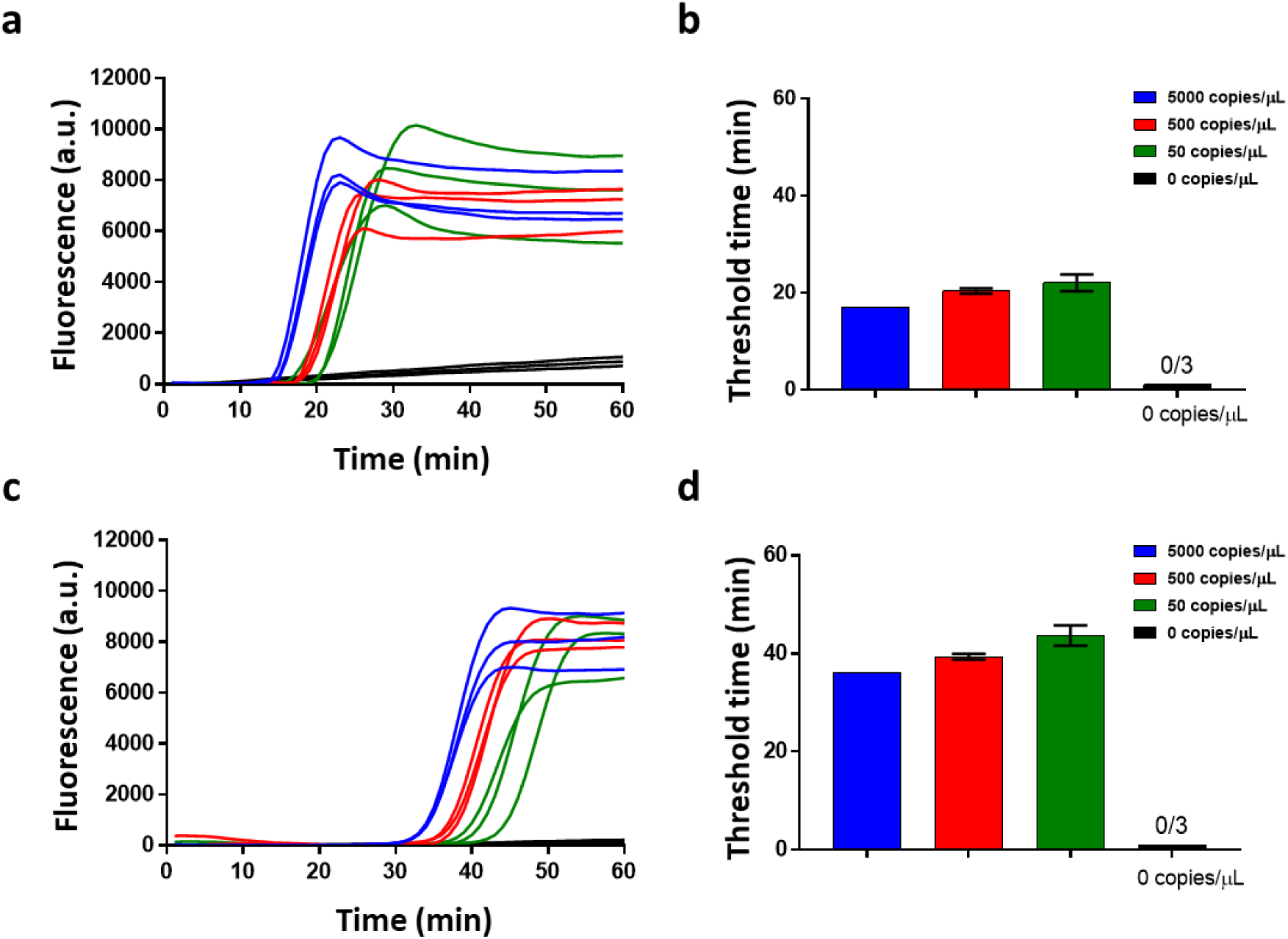
Characterization of SARS-CoV-2 virus in Viral Transport Media (VTM). **(*a-b*)** Raw fluorescence data and amplification threshold times (for 3 replicates of data) for viral detection in a 16 μL reaction with 12.5% spiked VTM sample in each reaction. **(*c-d*)** Raw fluorescence data and amplification threshold times (for 3 replicates of data) for viral detection in a 16uL reaction with 50% spiked VTM per reaction. Thermal lysis at 95°C was conducted for 1 min. of the virus in VTM sample before addition of RT-LAMP reagents for the final reaction. The bar graphs show mean and standard deviation.

### Assay characterization in simulated clinical workflow with Nasopharyngeal swab

To evaluate the performance of our detection assay in the current clinical workflow, we developed a protocol with simulated nasopharyngeal/nasal swab samples as shown in **Figure 6a**. Commercial swabs (Puritan sterile polyester tipped applicators, 25-800D 50) were introducing into purchased nasal solution spiked with known virus concentrations. Next, the swab was transferred to VTM and gently agitated in the solution to transfer the viruses from the swab into the VTM. The swab was thereafter discarded and aliquots from the VTM were taken to perform thermal lysis at 95°C for 1 min. Finally, RT-LAMP reagents were added, and the final reaction was performed at 65°C for 60 min. We transferred the mock swabs to 100 μL and 500 μL of VTM to evaluate VTM volume effects on swab viral load transfer efficiency. For each condition above, we performed limit of detection tests with 12.5% and 50% VTM per reaction. The amplification curves and threshold time bar graphs for reactions with 100 μL and 500 μL VTM are shown in **Figure S2a-d** and **Figure 6b-e**, respectively.

**Fig. 6.**
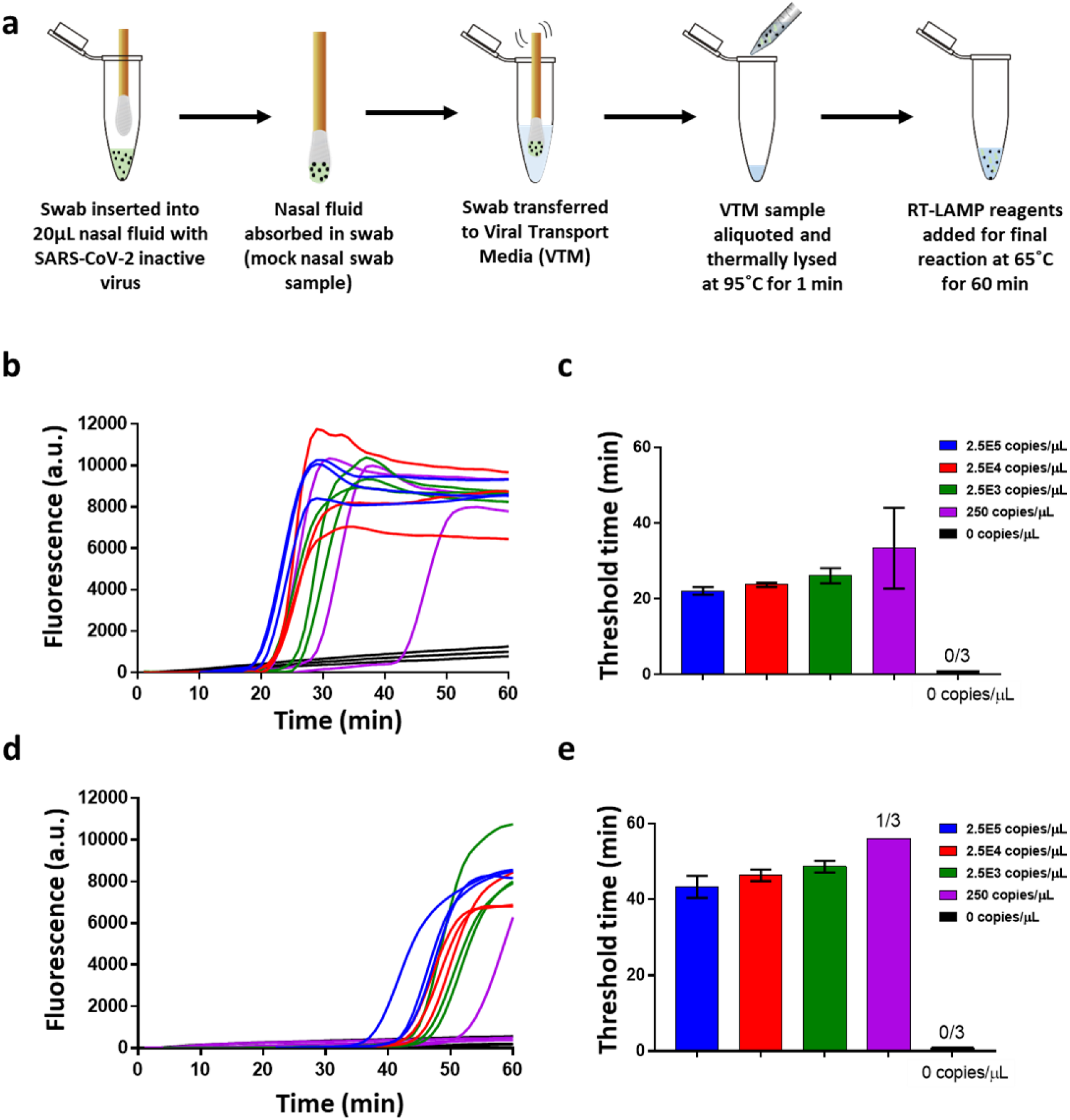
Characterization of SARS-CoV-2 virus in mock nasopharyngeal swab samples transported in Viral Transport Media. **(*a*)** Process flow schematic of viral detection from a mock nasopharyngeal swab. A swab is inserted into a tube with virus-spiked nasal fluid and absorbs the fluid. Post vigorously mixing the swab in 100 or 500uL of VTM, an aliquot of the VTM sample is then thermally lysed at 95C for 1 min. The RT-LAMP reagents are added to the lysed viral sample and the reaction is conducted at 65C for 60 min. **(*b-c*)** Raw fluorescence data and amplification threshold times (for 3 replicates of data) for viral detection in a 16 μL reaction with 12.5% VTM sample per reaction from a 500uL VTM sample. **(*d-e*)** Raw fluorescence data and amplification threshold times (for 3 replicates of data) for viral detection in a 16 μL reaction with 50% VTM per reaction from a 500 μL VTM sample.

For 100μl VTM, the limit of detection remained 2.5E4 copies/μL of virus in the starting nasal fluid for both 12.5% (**Figure S2a-b**) and 50% (**Figure S2c-d**) VTM in reaction, even though 50% VTM reactions showed delayed amplifications as previously observed. The above detection limit in nasal fluid amounts to 5E3 copies/μL of virus in VTM after swab transfer. This is two orders of magnitude greater than the 50 copies/μL detection limit of viruses directly spiked in VTM and indicates inefficient viral transfer from swab to 100 μL VTM solution, which is present in all swab-based sample collection processes. This inefficiency likely arises due to inadequate adsorption of viruses into the swab, and subsequent inadequate release of the viruses into the VTM. The low interfacial contact area between the swab and the VTM due to VTM volume could also play a role in the poor release of the viruses into the VTM.

For 500ul VTM with 12.5% (**Figure 6b-c**) and 50% (**Figure 6d-e**) VTM in reaction, the limit of detection improved to 250 copies/μL and 2.5E3 copies/ul of virus in nasal fluid, respectively. The above detection limits in nasal fluid amount to 10 copies/μL and 100 copies/μL of virus in VTM after swab transfer which are comparable to the control experiments where viruses were directly spiked in VTM. This indicates efficient viral release in 500 μL VTM with transfer efficiency close to 100%. Three swab replicates were performed for each condition. Our RT-LAMP assay process flow starting from a patient sample and its comparison to RT-PCR is shown in **Figure S3**, highlighting the fact that a viral extraction step is not required in our assay.

### SARS-CoV-2 detection directly from VTM in handheld reader and additively manufactured cartridges

Finally, we demonstrate detection of SARS-CoV-2 viruses in VTM using a portable handheld reader that could allow near-patient testing. In these tests, isothermal RT-LAMP reactions were performed on additively manufactured polymer cartridges where the samples were loaded manually and without the use of pumps. The portable handheld reader included heating elements and optics necessary for performing and recording the reaction, respectively. **Figure 7a** shows an exploded view of the handheld reader, detailing the components of the reader and **Figure 7b** shows an optical image of the reader with inserted cartridge. RT-LAMP reagents and virus-spiked VTM sample were loaded using syringes through luer-lock compatible inlet ports. The cartridge had 3D mixing features for improved mixing of the virus in VTM sample with the amplification reagents. **Figure 7c** shows the top and bottom plane of the serpentine mixing channels and zoomed-in images of the amplification and diagnostic regions with 6 pie-shaped amplification chambers. After mixing in the serpentine channels, the final reaction mix was flowed into the amplification chambers. The amplification reservoirs were specifically designed to allow uniform filling of all the chambers as can be seen in **Video S1** (Supplementary information). After the amplification chambers were completely loaded, they were sealed with biocompatible adhesive and the cartridge was inserted into the reader for the final reaction. The integrated heater was set at 65°C and a smartphone was used to perform the imaging. **Figure 8a-b** shows baseline-subtracted real-time fluorescence images of amplification on the cartridge for 5000 copies/μL of virus in VTM and negative control (VTM only). Video S2 and S3 (Supplementary information) show time stamped videos of amplification on the cartridge for 5000 copies/μL of virus in VTM and negative control (VTM only), respectively. The fluorescence images were analyzed using Image J software and the mean fluorescence intensity of the 6 amplification chambers were plotted over time in **Figure 8c**. A positive detection can be seen in as low as 30 minutes from the start of the reaction. **Figure 8d** shows the endpoint fluorescence images at 40 minutes of three reaction replicates for 5000 copies/μL of virus in VTM and negative control. The mean fluorescence intensity of these endpoint results is shown in **Figure 8e** clearly showing the difference between the two groups. Together, these results demonstrate the ability of our platform to provide rapid SARS-CoV-2 detection results from unprocessed VTM samples in a portable hand-held reader.

**Fig. 7.**
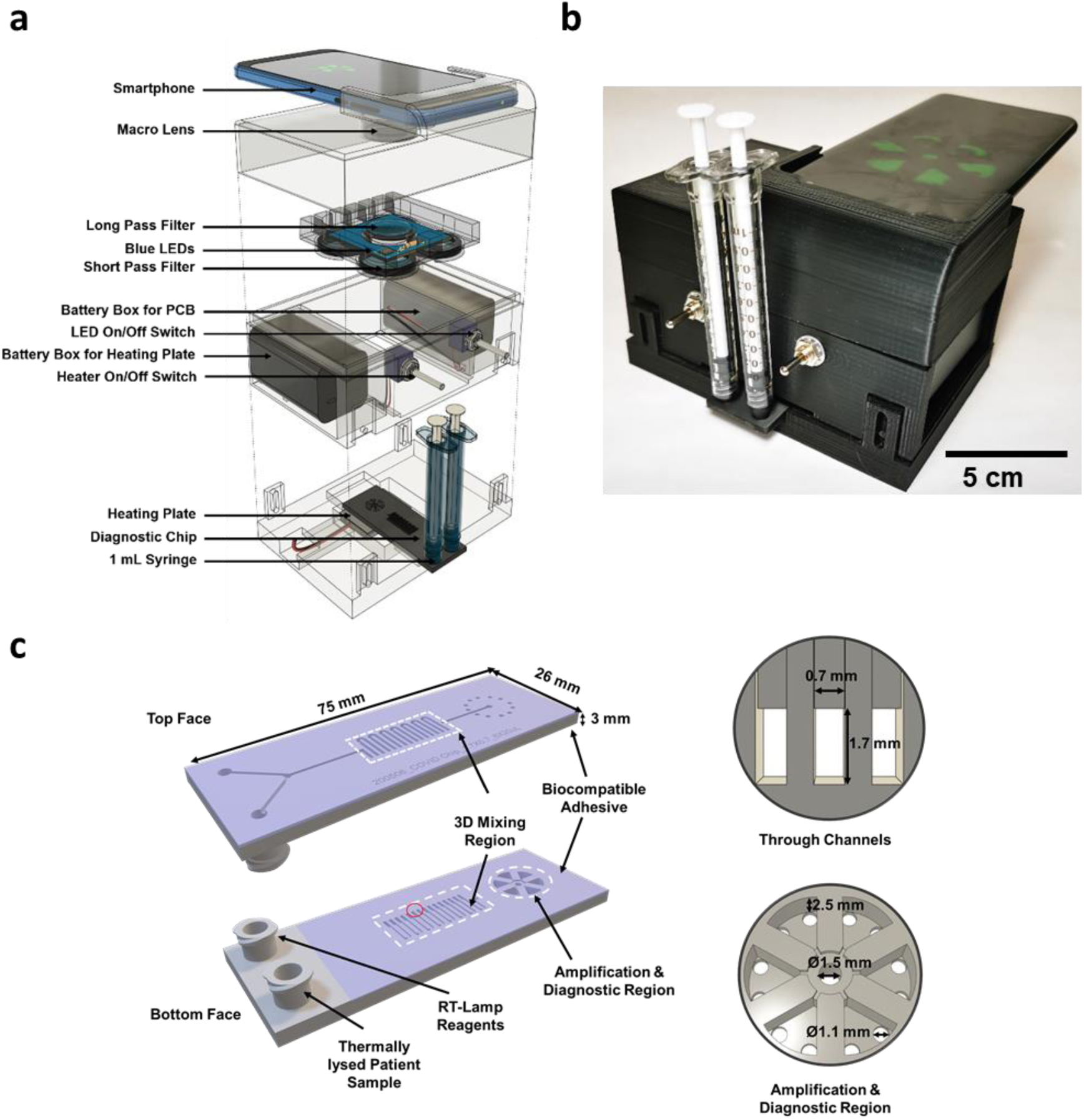
Handheld point of care instrument and additively manufactured cartridge. **(*a*)** Schematic of the handheld point of care instrument showing components in an exploded view. A smartphone images the cartridge, while isothermal heating and illumination are battery powered. Optical components integrated with the instrument match the excitation and emission characteristics of the fluorescent signal. **(*b*)** Photograph of the instrument used for rapid detection. **(*c*)** Diagram of the diagnostic cartridge used for rapid detection of SARS-CoV-2 in VTM. Two inlets mate with syringes that inject either RT-Lamp Reagents or Thermally Lysed Patient Sample into a 3D serpentine mixing region before filling the amplification and diagnostic region. Device dimensions are highlighted for the 3D mixer and amplification and diagnostic region.

**Fig. 8.**
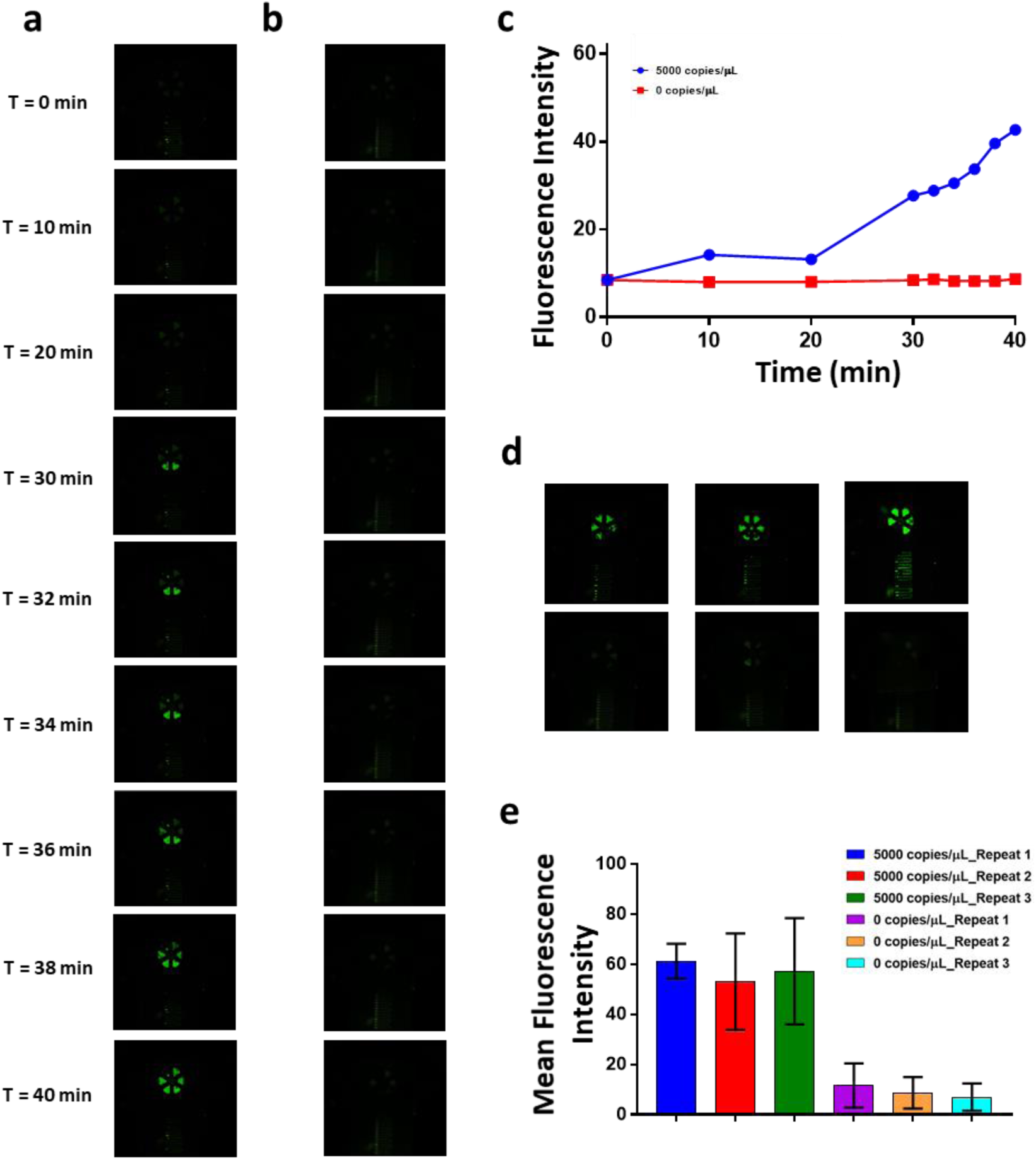
Rapid Detection of SARS-CoV-2 in Viral Transport Media in an additively manufactured cartridge and handheld point of care instrument. **(*a*)** Baseline-subtracted fluorescence images of real-time RT-LAMP on the additively manufactured amplification chip at different time points showing the amplification of 5000 copies/μL of SARS-CoV-2 inactive virus in VTM. **(*b*)** Baseline-subtracted fluorescence images of real-time RT-LAMP on the additively manufactured amplification chip at different time points showing the amplification of VTM only negative control. **(*c*)** Baseline subtracted mean fluorescent intensity over time for on-chip amplification detection. **(*d*)** Amplification chip endpoint fluorescence images at time = 40 min. (for 3 chip replicates) of 5000 copies/μL of SARS-CoV-2 inactive virus in VTM (top) and, amplification chip endpoint fluorescence images at time = 40 min. of VTM only negative control (bottom). **(*e*)** Mean fluorescent intensity at time = 40 min. (for 3 chip replicates) for 5000 copies/μL of SARS-CoV-2 inactive virus in VTM and VTM only negative control. The bar graphs show mean and standard deviation of the fluorescence intensity of the 6 pools in a single chip replicate.

### Quantitative model of resources required for our RT-LAMP assay compared to standard RT-PCR test to perform testing at scale

We developed a model to quantify the resources required to scale up the RT-LAMP assay and compare it to the conventional RT-PCR test. For each test, we considered three scenarios in which the number of patient samples are 80, 800, or 8000. **Table 1** shows the key results for cost and time required. The supplementary information shows additional data (**Figure S4**) and the complete resources model for the different scenarios (**Table S2**).

**Table 1.** Cost and personnel required to perform the analysis of 80, 800, and 8000 samples.

| **Number of tests** | **80** |  | **800** |  | **8000** |  |
| --- | --- | --- | --- | --- | --- | --- |
| **Technique** | RT-LAMP | RT-PCR | RT-LAMP | RT-PCR | RT-LAMP | RT-PCR |
| **Cost per test excluding labor (\$)** | 51.7 | 61.4 | 10.4 | 16.4 | 6.6 | 11.9 |
| **Cost per test including labor (\$)** | 76.2 | 92.4 | 34.9 | 47.4 | 31.1 | 42.9 |
| **Consumables cost per test (\$)** | 49.4 | 59.1 | 9.1 | 15.3 | 6.3 | 11.6 |
| **Total time (h)** | 5.5 | 6.9 | 10.9 | 13.8 | 54.6 | 68.9 |
| **Number of People** | 6 |  | 30 |  | 60 |  |
| **Number of thermocyclers** | 1 |  | 5 |  | 10 |  |

Different scenarios require different quantities of labor and laboratory resources. Many laboratory instruments are capable of 96 parallel tests, and thermocycler loaded with 96 tests performing the CDC RT-PCR assay would have 80 patient samples plus eight positive controls and eight negative controls^32^. Thus, we select 80 as the smallest increment of patient samples and multiples of 80 for scale up. The model accounts for the time and cost to process 16 control samples for every 80 patient samples. For the 80 patient test scenario, we assume a work force of 6 people and use of one thermal cycler, for 800 patient tests we assume 30 people and 5 thermal cyclers, and for 8000 patient tests we assume 60 people and 10 thermal cyclers. Other laboratory infrastructure is required as well including refrigerated sample storage, biohazard waste management along with sufficient space to work. The cost modeling includes the costs of the reagents and disposable supplies such as swabs, pipettes, and vials, the cost of labor at $1 / minute, and the time averaged cost of using laboratory instruments calculated as the instrument cost divided over 10,000 hours of useful lifetime. The model does not account for the cost of space as this could vary widely. The modeling also assigns an expected time required for each step in the process. All of the costs are tracked either to the CDC test for RT-PCR^32^ or the steps described in the present study for RT-LAMP. See **Table S2** for detailed breakdown of all of the costs and time required for each step.

The modeling shows that RT-LAMP is faster and less costly than RT-PCR, and these advantages can be linked to two key differences between the assays. First, the amplification time for RT-LAMP is about half of that for RT-PCR. RT-PCR requires time to progress through successive thermal cycles while RT-LAMP requires only one isothermal step. Second, RT-LAMP does not require a separate step for RNA extraction, saving time and reducing the cost of consumables. While the elimination of the RNA extraction kit has advantages, a disadvantage is that additional controls are required to account for the presence of the swabs. The swabs could be eliminated if other type of specimens such as saliva could be used for the assay. For example, a recent report shows direct RT-PCR with patient nasopharyngeal samples without nucleic acid extraction^35^.

## Discussion

The current gold standard method for the detection of SARS-CoV-2 virus is a PCR-based analysis, which requires laboratory-based protocols for viral isolation, lysis, and removal of inhibiting materials. Likewise, PCR requires precise thermal cycles, typically controlled using a thermocycler instrument, to amplify the RNA sequences. These requirements constrain the ways that diagnostic testing can be delivered and limit access to testing in certain situations and regions of the world. There is an urgent need for accessible, low-cost, and portable platforms that can provide fast and accurate diagnosis at the point-of-use.

In this paper, we present an isothermal RT-LAMP based assay for rapid detection of SARS-CoV-2 and demonstrate the feasibility of a point-of-use approach. We show the assay development and demonstrate 50 copies/μL limit of detection with genomic RNA and inactive whole viruses. The assay can robustly detect the virus directly from nasal fluid and viral transport media (VTM). There was is loss of sensitivity in the VTM and the detection limit remained constant at 50 copies/μL. In a simulated clinical workflow with nasal swab in VTM, efficient viral transfer from swab to VTM occurrs for 500 μL of VTM volume but not for 100 μL, perhaps due to lower interfacial contact area between the VTM solution and the swab for the 100 μL scenario. Finally, we demonstrate rapid (< 40 minute) detection of virus directly from VTM using a portable hand-held reader and additively manufactured cartridges.

There have been a few recent reports on the development of RT-LAMP-based assays to detect SARS-CoV-2 from nucleic acids. These studies have highlighted the minimal instrumentation required^36^, LOD around 120 copies/reaction^37^, and no cross-reactivity against MERS, BtCoV, MHV, other strains (human coronavirus strains 229E, NL63, HKU1, or OC43)^38^, or PEDV, TGE, PDCoV, and IBV^36^. To the best of our knowledge, no previous study has shown RT-LAMP based detection from SARS-CoV-2 virus. Similarly, no previous study has shown on-chip detection of SARS-CoV-2 using a handheld instrument.

RT-LAMP has key advantages for speed and cost compared to RT-PCR. Importantly, a batch of 80 patient samples can be tested about 1 hour faster using RT-LAMP compared to RT-PCR and is around 20% lower cost. When scaling up to larger batches, 800 or 8000 patient samples, the speed and cost advantages increase significantly. The use of automation to reduce the labor content would reduce the time and cost of both approaches, however this would increase the advantages that RT-LAMP has over RT-PCR, because the specific sources of time and cost reduction for RT-LAMP would account for a larger portion of the overall time and cost.

A point-of-use system testing a single patient sample provides even larger advantages over laboratory-based RT-PCR. The a single patient test in a point-of-use system using RT-LAMP allows collection and diagnostic to be performed in about 50 minutes at the point of sample collection, while a single patient test using RT-PCR requires transport time to a laboratory plus more than 90 minutes in the laboratory for the complete diagnostic. The point-of-use system also has lower capital costs at around $1000 for the entire system. The availability of a rapid, inexpensive, and easy to use system for detection of SARS-CoV-2 could increase access to testing in many situations and communities.

## Materials and Methods

### SARS-CoV-2 Genomic RNA and viruses

Genomic RNA for SARS-Related Coronavirus 2 (Isolate USA-WA1/2020), NR-52285, was obtained from BEI Resources. These genomic RNA vials were stored at -80°C and stock volumes were either used directly for experimentation or diluted to the correct concentration in TE Buffer. For experiments using virus, Heat Inactivated SARS-Related Coronavirus 2, NR-52286, was obtained through BEI resources. These stocks were aliquoted and stored at -80°C. Either stock volumes were used for direct experimentation or diluted in TE Buffer or Viral Transport Media to the correct concentrations.

### Primer sequences and primer validation reactions

Primer sequences for the RT-LAMP reactions were synthesized by Integrated DNA Technologies (IDT) and are listed in Table S1 (Supplementary information). Primerexplorerv4 (https://primerexplorer.jp/e/) was used to design all sets of RT-LAMP primers for SARS-CoV-2 RNA and virus. The sequence for SARS-CoV-2 virus was obtained from the NCBI database (Genebank number - MN988713.1).

In total 12 sets of primers (3 sets of primers each for 4 different gene targets) were tested with SARS-CoV-2 genomic RNA as template to determine the best primer set for each gene that detected 500 copies of RNA/μL with the lowest threshold time. Primer set 3 for Gene orf1a, primer set 2 for Gene S, primer set 2 for Gene orf 8, and primer set 1 for Gene N were selected. Thereafter, RT-LAMP assays with the four selected primer sets were conducted on ten-fold serially diluted RNA to determine the detection range. The Gene N targeting primer set (Primer set 1) showed detection of 50 genomic copies/μL of SARS-CoV-2 genomic RNA as the limit and was therefore selected as the working primer set used in the downstream RT-LAMP assays.

### Genomic RNA and virus in Buffer detection in RT-LAMP reactions

The following components comprised the RT-LAMP assay: 4 mM of MgSO4 (New England Biolabs), 1× final concentration of the isothermal amplification buffer (New England Biolabs), 1.025 mM each of deoxy-ribonucleoside triphosphates (dNTPs), and 0.29 M of Betaine (Sigma-Aldrich). Individual stock components were stored according to the manufactural instructions and a final mix including all of the components was freshly created prior to each reaction. Along with the buffer components, a primer mix consisting of 0.15 μM of F3 and B3, 1.17 μM FIP and BIP, and 0.59 μM of LoopF and LoopB was added to the reaction. Finally, 0.47U/μL BST 2.0 WarmStart DNA Polymerase (New England Bioloabs), 0.3 U/μL WarmStart Reverse Transcriptase (New England Biolabs), 1 mg/mL BSA (New England Biolabs), and 0.735x EvaGreen (Biotium) was included in the reaction. EvaGreen dye is a double-stranded DNA intercalating dye. After addition of the template, the final volume of the reaction was 16μL. All reactions with genomic RNA template in TE Buffer included 2 μL of template to make the final reaction volume of 16 μL.

All of the off-chip RT-LAMP assays were carried out in 0.2 mL PCR reaction tubes in an Eppendorf Mastercycler® realplex Real-Time PCR System at 65°C for 60 min. Fluorescence data was recorded every 1 min. after each cycle of the reaction. Three repeats were done for each reaction. Reactions done with heat-inactivated viruses included a thermal lysis step. First serially diluted in TE Buffer, viral samples were then thermally lysed in a heater at 95°C for 1 min. prior to their addition into the final reaction mix.

All the RT-LAMP reactions consisted of non-template negative controls that were included in all the datasets.

### Detection of virus spiked in nasal fluid

Nasal fluid was commercially obtained from Innovative Research and it was confirmed that the fluid samples were obtained prior to the COVID19 pandemic. Serially diluted SARS-CoV-2 heat-inactivated viruses in TE Buffer were spiked directly into nasal fluid such that the viral sample concentration in nasal fluid ranged from 50 to 0.005 PFU/μL (5E5 to 50 copies/μL). The virus in Nasal fluid sample were then thermally lysed at 95°C for 1 min. prior to adding the RT-LAMP reagents mix for a total reaction volume of 16 μL. The sample volumes were varied such that the spiked nasal fluid sample volume 12.5%, 25%, or 50% of the total reaction volume. One reaction was conducted in the same format in which total reaction volume was 96 μL of which 48 μL was the sample volume. In all of these reactions, the concentrations of all other reaction’s components were maintained as mentioned above for the 16 μL reaction.

### Detection of virus in Viral Transport Media

Moreover, reactions were done with heat inactivated viruses in Viral Transport Media (VTM). CDC Compliant Viral Transport Media was obtained from Redoxica (VTM-500ML), aliquoted, and stored in 4°C away from direct light. Viruses were serially diluted in VTM to starting sample concentrations ranging between 0.5 and 0.005 PFU/μL (5000 copies/μL to 50 copies/μL). Then, the samples were thermally lysed at 95°C for 1 min. prior to adding the RT-LAMP reagents mix for a total reaction volume of 16 μL. Two different sample volumes were tested in which the virus in VTM sample was either 12.5% (2 μL sample) or 50% (8 μL sample) of the total reaction volume. In these reactions, the concentrations of the buffer, primer, polymerase, and other reaction components were kept constant as mentioned above within the 16 μL reaction.

### Detection of virus in nasal fluid collected on nasopharyngeal swabs

CDC approved Nasopharyngeal swabs (Sterile Polyester Tipped Applicators) were commercially obtained from Fisher Scientific. As described above, serially diluted SARS-CoV-2 heat-inactivated viruses in VTM were spiked directly into nasal fluid such that the viral sample concentration in nasal fluid ranged from 25 to 0.0025 PFU/μL (2.5E5 to 250 copies/μL). Spiked nasal fluid (20 μL) was first absorbed by a nasopharyngeal swab and then transferred into 100 μL of VTM. The swab was mixed in the VTM for 30 s. to 1 min. for viral transfer from the nasopharyngeal swab. After removing the swab, the VTM sample was distributed into sample aliquots and thermally lysed at 95°C for 1 min. Finally, the rest of the reagents for the RT-LAMP reaction was added to the sample for a total reaction volume of 16 μL.

### Off Chip amplification data analysis

The off-chip RT-LAMP fluorescence curves and amplification threshold bar graphs were analyzed using a MATLAB script and plotted using GraphPad Prism 7. For each curve, the threshold time taken as the time required for each curve to reach 20% of the total intensity. The amplification threshold bar graphs are show a mean and standard deviation of 3 samples.

### Additively manufactured cartridge fabrication

Figure 8(c) shows the disposable polymer cartridge developed for the rapid detection of SARS-CoV-2 in VTM. The 3D design consists of microfluidic channels on both front and back sides, connected by 1.7 mm X 0.7 mm2 through holes at the end of each serpentine microchannel. The chip was designed and additively manufactured as a single component to complete three functions on-chip. First, thermally lysed patient sample and RT-LAMP reagents are injected through the female luer lock connectors from two separate syringes without the use of microfluidic pumps. Each access port is directly connected to the continuous 3D flow pathway by a Y-shaped inlet region. Then the sample flows through the 3D micromixer region, where the flow takes a vertical turn from one face to the other face between each horizontal U-turn. There are seven serpentine channels on the top face and eight on the bottom face with each serpentine microchannel being 0.7 mm wide, 0.4 mm deep, and 8 mm long. The alternating horizontal and vertical U-turns enhance mixing and allow for dense packing of the mixing structure. Finally, the fluid flows into six reservoirs that radially surround the flow channel furcation. These detection reservoirs located at the end of the chip are designed to contain a volume of ∼20 μL per chamber. The amplification chambers have a 0.5 mm thick wall and two 1.1 mm diameter outlet holes to remove excess air during filling.

The cartridge was fabricated from RPU on a Carbon M2 printer using standard process settings, washed, and then cured. The print orientation for this part was the bottom face of the chip attached to build tray, and luer lock fluid ports facing upwards. After fabrication, the cartridge was washed again in water and dried using pressurized air. The front and back sides of the cartridge were covered with transparent biocompatible tape (ARSeal 90880, Adhesive Research) to seal the chip. The transparent tape allows for visual inspection during filling and optical imaging during detection. Following tape application, two holes were made using a needle in the tape for each reservoir; these holes serve as air outlets during filling.

### Cradle fabrication and smartphone-based fluorescence imaging

The microfluidic diagnostic cartridge mates with an instrument shown in Fig 7 and 7b. We used a smartphone (Huawei P30 Pro, Huawei) to detect the fluorescence emission from on-chip LAMP assays. The instrument comprises four main parts to support optical, electrical, and heating components. The top part of the instrument holds the smartphone and aligns its camera with a macro lens (12.5X, Techo-Lens-01, Techo). The macro lens enables close-up imaging (∼50 mm imaging distance) of the chip. The second component, PCB & filter holder, is equipped with a long pass filter (525 nm, 84-744, Edmund Optics), which allows only the emission light from the EvaGreen dye to reach the camera. A printed circuit board (PCB) is aligned with this long-pass filter and controls the illumination of the device. A total of eight light-emitting diodes (LED) (λpeak = 485 nm, XPEBBL, Cree) are mounted on the PCB in a circle to provide uniform illumination over the diagnostics area. Four short pass filters (490 nm, 490SP RapidEdge, Omega Filters) covering each pair of LEDs are mounted on top of PCB to excite the EvaGreen dye. The third component, namely, the main body of the cradle, carries two separate on-off switches and battery boxes to control the PCB and a heater. Finally, a self-regulating positive temperature coefficient (PTC) heater (12 V-80 °C, Uxcell) is located below the cartridge. A locking mechanism connects the main and bottom components opens and grants access to the chip to be inserted into the cradle. When inserted, only the diagnostics area of the chip is in contact with the heating plate that keeps the temperature at 65 °C during the amplification period.

The smartphone took photos of the chip at 10-minute intervals in the first 30 minutes of assays, then at 2-minute intervals to capture more amplification data points for another 10 minutes. The imaging settings of the smartphone were ISO = 500 and exposure time = 0.5 s.

### Chip loading of viral sample and RT-LAMP assay components

For on chip experiments with viruses in VTM, the viruses were spiked directly in VTM such that their concentration in the sample were 0.5 PFU/μL (5000 copies/μL). The sample was loaded into a 1 mL syringe. The RT-LAMP reagents were prepared off-chip and then loaded into a 5mL syringe. All concentrations of reagents were maintained as in a 16 μL reaction. Both syringes were attached to the chip and the sample was loaded on chip without the use of a syringe pump. Once the sample and reaction reagents were loaded into the chip, the holes underneath the pie pools were sealed with a second double-sided adhesive layer to prevent leakage or evaporation during RT-LAMP incubation. The chip was placed into a portable cradle and clamped down with magnetic strips, such that the pie pools made good contact with the PTC heater throughout the RT-LAMP incubation step. The incubation occurred at 65°C for 60 minute with real-time monitoring.

### Chip Image and data analysis

Fluorescence images were recorded with IP Webcam in smartphones and were saved in JPG format from which fluorescence intensity and baseline fluorescence was analyzed on Image J. The fluorescence intensity from all 6 pie pools were measured, averaged, and plotted on GraphPad Prism 7.

## Acknowledgments

The following reagent was deposited by the Centers for Disease Control and Prevention and obtained through BEI Resources, NIAID, NIH: (i) Genomic RNA from SARS-Related Coronavirus 2, Isolate USA-WA1/2020, (ii) NR-52285. SARS-Related Coronavirus 2, Isolate USA-WA1/2020, Heat Inactivated, NR-52286.

Authors thank the staff at the Holonyak Micro and Nanotechnology Laboratory at UIUC for facilitating the research and the funding from University of Illinois. RB, EV acknowledge support of NIH R21 AI146865A, and to also support AG. AM was partially supported by a cooperative agreement with Purdue University and the Agricultural Research Service of the United States Department of Agriculture, AG Sub Purdue 8000074077 to RB. The authors gratefully acknowledge the support of cooperative agreement #D19AC00012 awarded by the Defense Advanced Research Projects Agency of the U.S. Department of Defense to WPK and RB to support MA, JB, and EV. F.S. was supported by National Science Foundation (NSF) under the grant number 1534126.

## Supplementary information

**Fig. S1.**
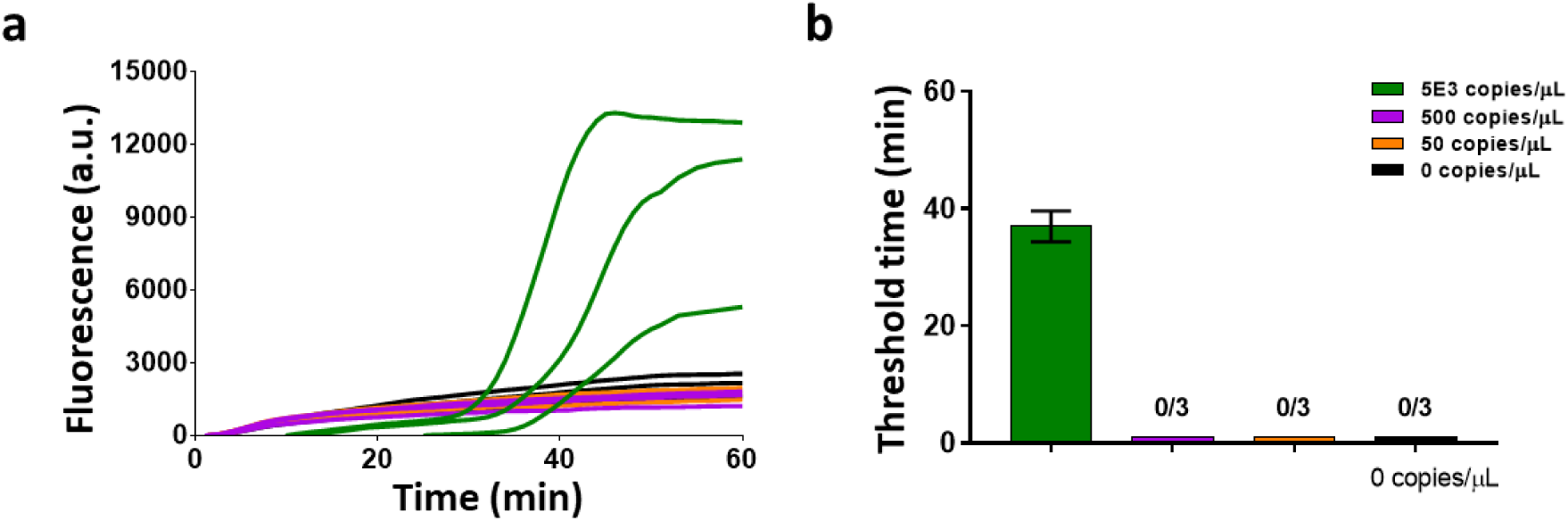
Characterization of SARS-CoV-2 virus in nasal fluid in a 96 μL reaction. **(*a-b*)** Raw fluorescence data and amplification threshold times (for 3 replicates of data) for viral detection in a 96 μL reaction with 50% nasal fluid per reaction. Thermal lysis at 95°C was conducted for 1 min. of the virus in nasal fluid sample before addition of RT-LAMP reagents for the final reaction. The bar graphs show mean and standard deviation. Fraction indicates number of replicates amplified.

**Fig. S2.**
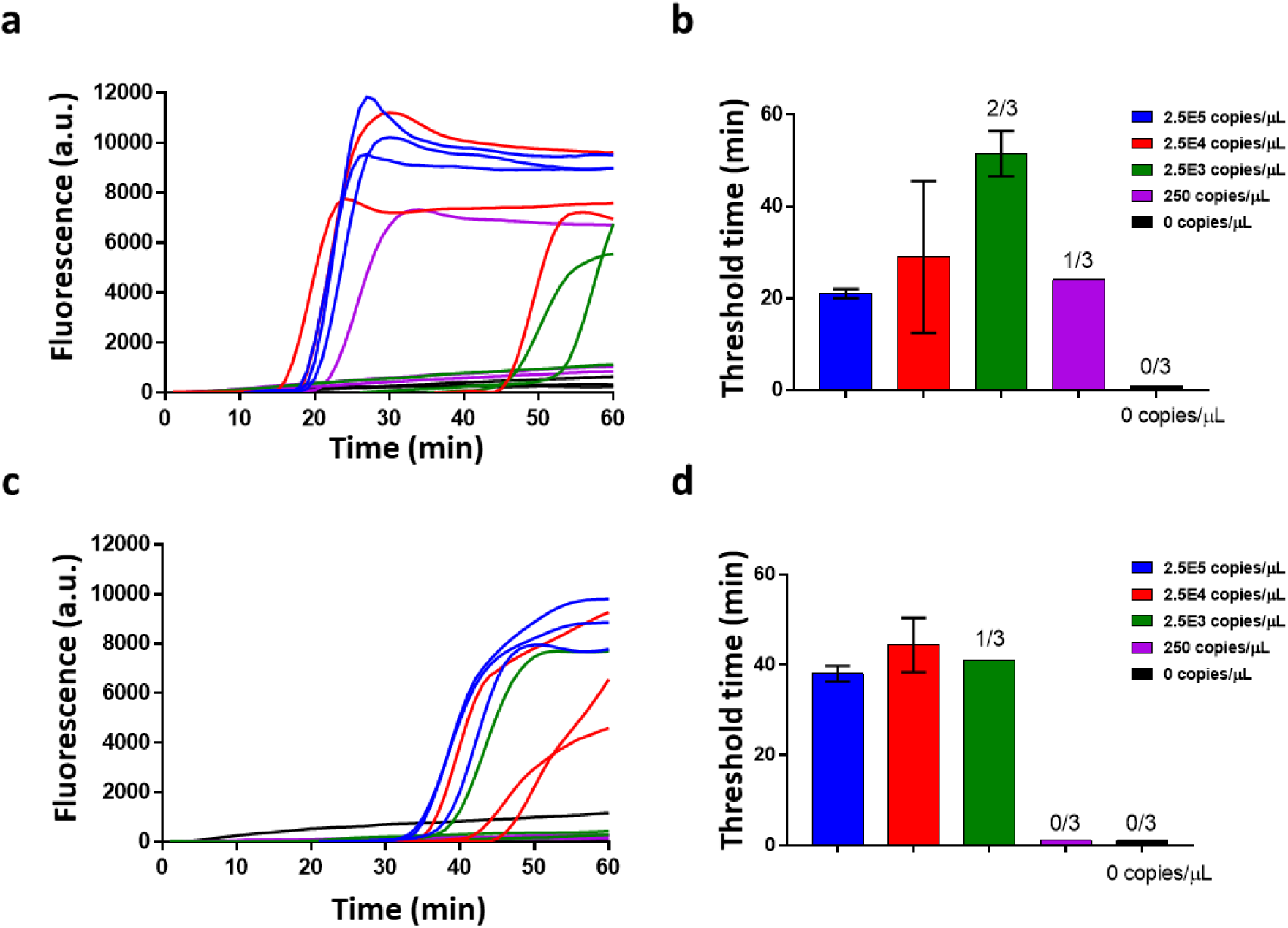
Characterization of SARS-CoV-2 virus in mock swab samples transported in 100 μL Viral Transport Media. **(*a-b*)** Raw fluorescence data and amplification threshold times (for 3 replicates of data) for viral detection in a 16 μL reaction with 12.5% VTM per reaction from a 100 μL VTM sample. (***c-d***) Raw fluorescence data and amplification threshold times (for 3 replicates of data) for viral detection in a 16 μL reaction with 12.5% VTM per reaction from a 100 μL VTM sample. The bar graphs show mean and standard deviation.

**Fig. S3.**
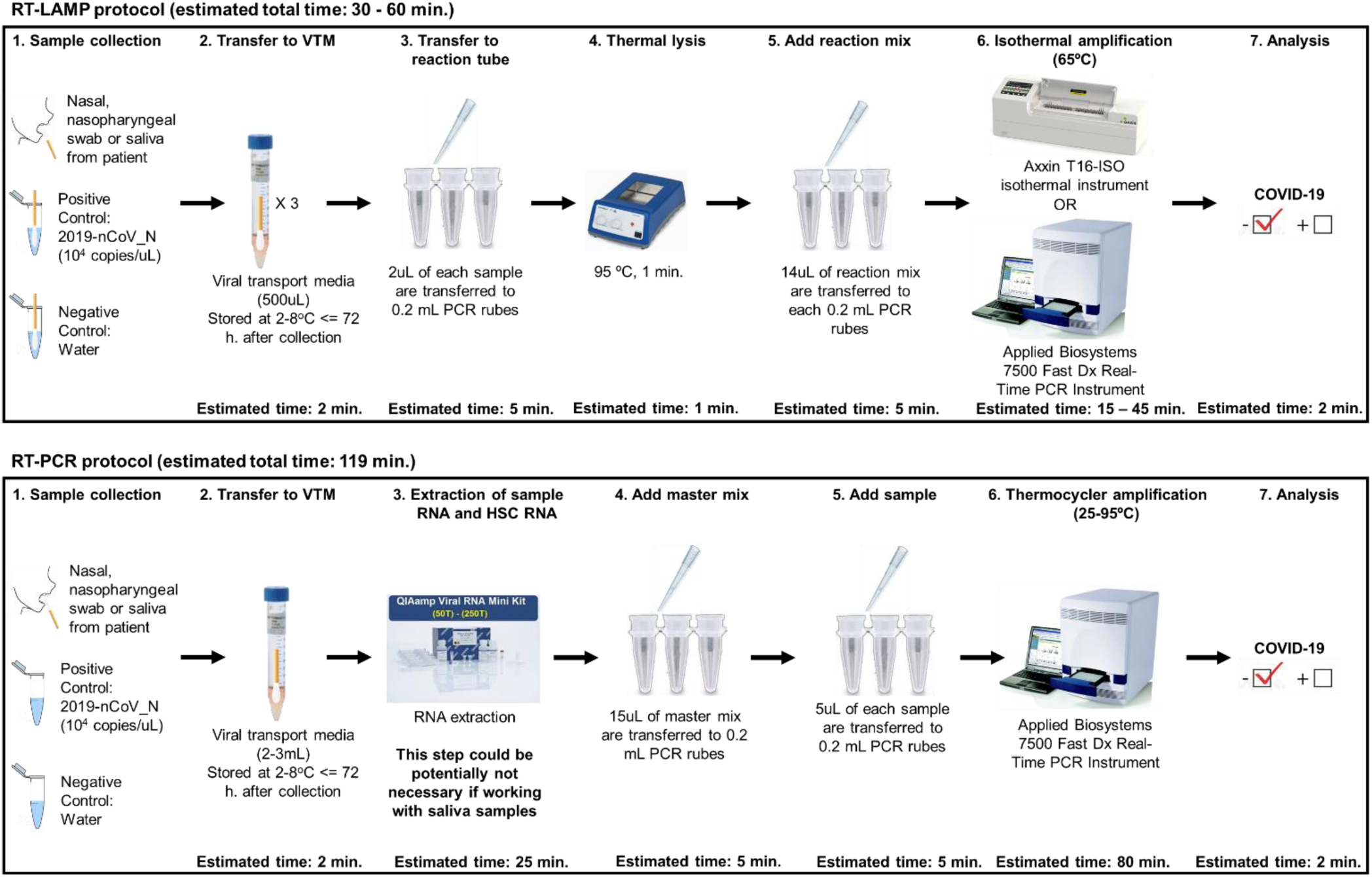
RT-LAMP assay process flow and comparison side by side with the RT-PCR assay process flow.

**Fig. S4:**
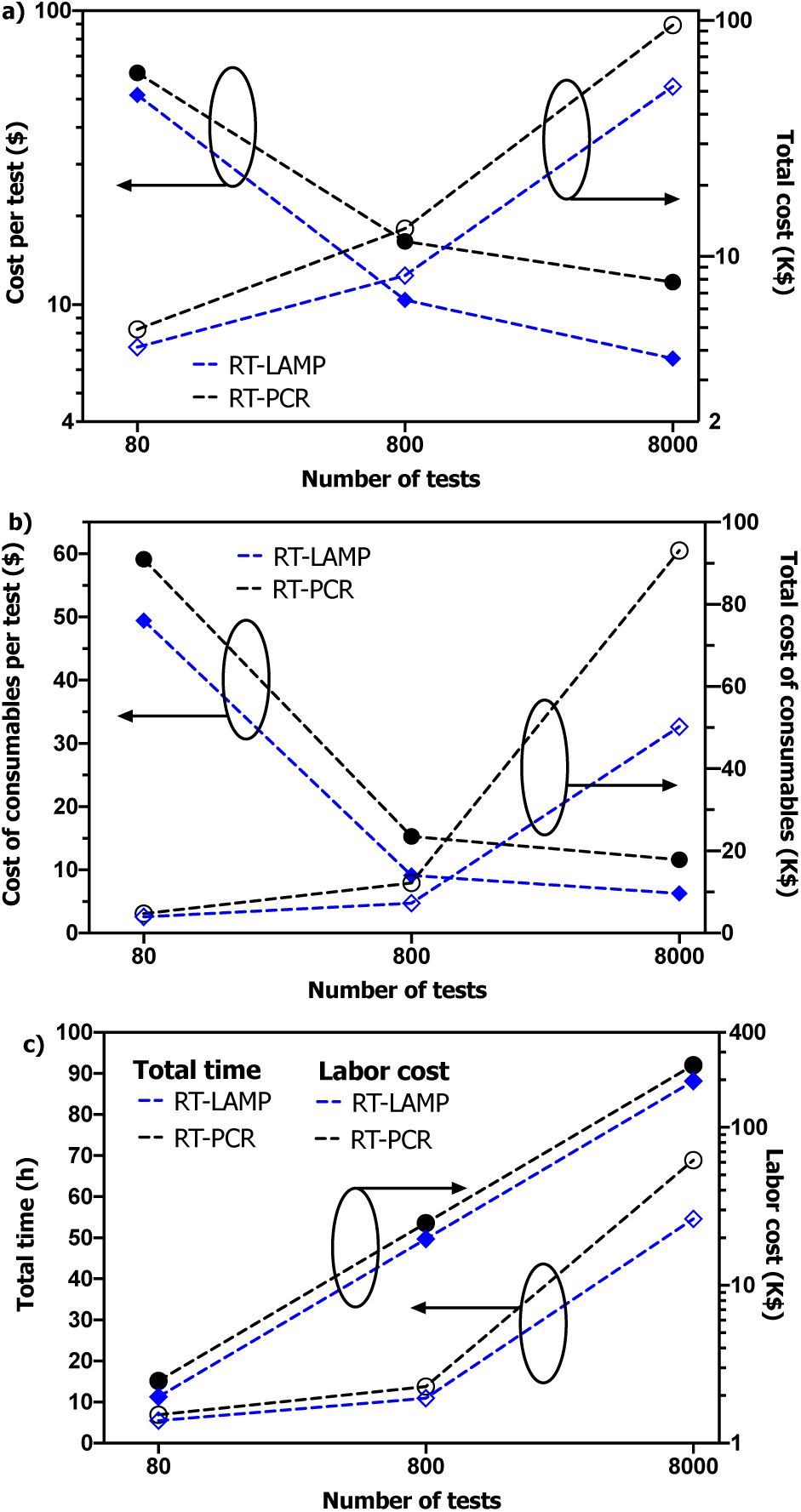
Resources modeling results. a) Cost per test and total cost (excluding labor) for the three scenarios considered; b) Total cost of consumables and Cost of consumables per test for the 3 models designed; c) Labor cost and Total time for the 3 models designed. The labor cost assumes a gross salary rate = $1/min.

**Table S1.**
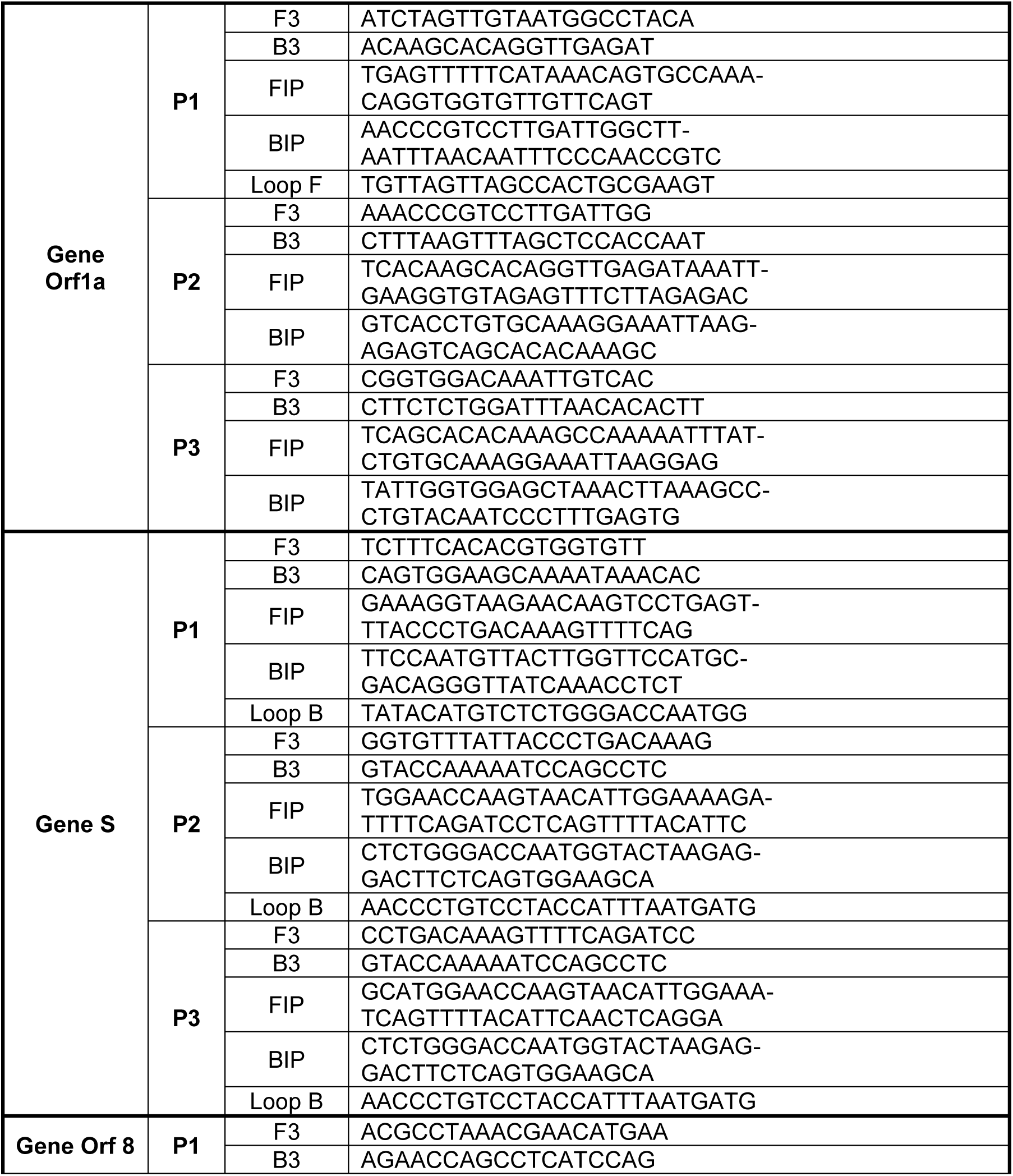
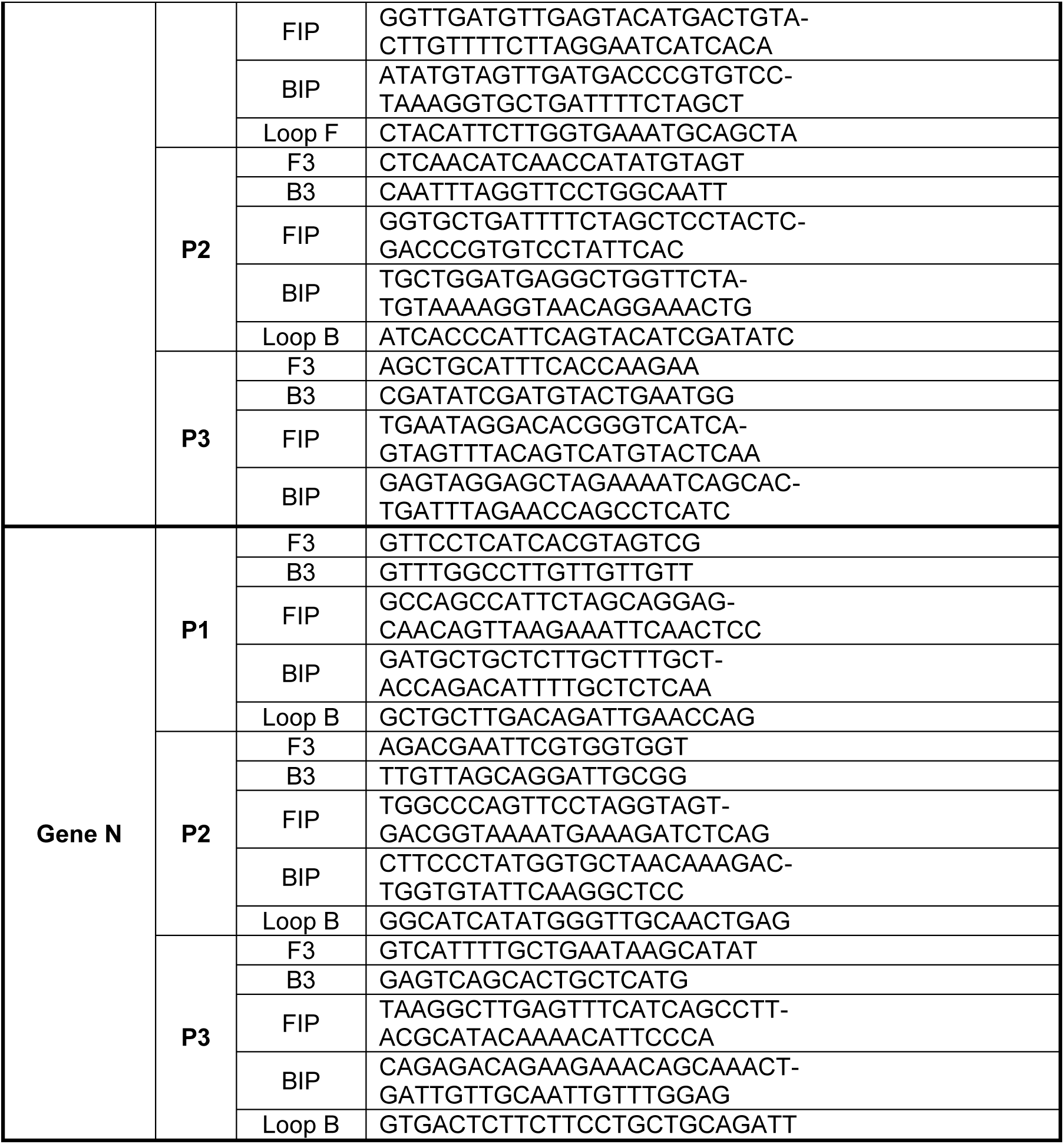
RT-LAMP primer sequences for 12 primer sets (3 primer sets each for 4 target genes)

**Table S2. Supply chain: RT-LAMP and RT-PCR assay**

See excel file.

**Video S1. Uniform filling of the amplification chambers.**

See video

**Video S2. Time stamped videos of amplification on the cartridge for 5000 copies/μL of virus in VTM.**

See video

**Video S3. Time stamped videos of amplification on the cartridge for negative control (VTM only).**

See video

